# *In vivo* Protective Efficacy of Emodin in Swiss Albino Mice Induced with Dalton Ascitic Lymphoma

**DOI:** 10.1101/2024.10.17.618958

**Authors:** Jagadish Kumar Suluvoy, Harish Babu Kolla, Jesse Joel, Aavany Balasubramanian

## Abstract

Lymphoma is a tumor that affects lymphoid tissues in the body. Treating lymphoma has become challenging because of the complexity of disease pathology, drug resistance mechanisms and side effects of existing chemo and radiation therapies. Treating cancers/tumors with plant based natural compounds is gaining interest recently because of their less toxicity profiles and efficiency in controlling the disease severity. Emodin is one such compound with such anti-cancer/tumor properties. It has immunosuppressive and anti-cancer properties through multiple ways. In this study, we have studied the therapeutic effect of emodin molecule in the DAL induced lymphoma, a well-established murine model to study and test the anti-lymphoma drugs. Our data has shown an outstanding therapeutic effect of emodin in controlling the lymphoma readouts in DAL induce Swiss Albino mice. These effects were studied in comparison with a standard drug molecule called methotrexate. Furthermore, the *in-silico* analysis has shown that emodin as a potential drug candidate for lymphoma based on the Lipinski’s rule of 5.

## Introduction

Lymphoma is a state of tumor characterized by the proliferation of lymphoid tissues in the body (Jiang et al., 2017). It is considered as a severe health burden globally with high mortality and morbidity rates (Chu et al., 2023). Clinically, lymphoma patients are being treated with chemotherapeutic drugs and radiation (Anand et al., 2024). These approaches have their own limitations like drug resistance, minimal efficacy and severe side effects or toxicity due to the nature of treatments. Thus, their limitations underscore the immediate requirement of novel and effective drugs or therapeutics to enhance the current lymphoma treatment with minimal side effects.

In this scenario, natural compounds from plant origin have gained attention for treating cancers because of their less toxicity and diverse biological activities ranging from anti-cancer/anti-tumor to immunomodulatory, anti-microbial and anti-inflammatory agents. Like and among the other plant based natural compounds, emodin compound has shown its promising anti-cancer properties. Emodin is a natural anthraquinone obtained from various plants such as Aloe vera, Chinese Goldthread and Rheum sp etc (Dong et al., 2020; Stompor-Goracy, 2021). It has been shown that the emodin induce apoptosis and further inhibit the cell proliferation in cancer cells (Zhang et al., 2020). This nature makes it a potential anti-cancer agent for the further investigation in lymphoma treatment.

Furthermore, use of murine lymphoma models such as Dalton’s Lymphoma (DAL)-induced models provides a valuable platform to study the disease pathogenesis, test and evaluate the anti-lymphoma drugs in laboratory settings (Debnath et al., 2018). DAL is a very well characterizes T-cell lymphoma model that mimics the pathology of human lymphoid malignancies. This platform offers an insight into the progression of tumor and study the therapeutic effect of various drugs before translating them into patient treatment. Moreover, previous studies have shown that the DAL can be effectively induced in the Swiss Albino mice and lead to the development of human like lymphoma pathological condition, thus serving as a better murine model to study lymphoma (Zhou et al., 2023).

In this study, we aim to study the *in vivo* therapeutic efficacy of emodin in the DAL-induced Swiss Albino mice through several gross pathological, hematological, biochemical, and histopathological readouts. We also investigated the drug-likeliness of emodin through *in-silico* resources. The findings highlight that the emodin treatment is efficient significantly controlling or improving various readouts of lymphoma in DAL induced Swiss Albino mice. The *in-silico* analysis also has shown that the emodin as a potential to treat lymphoma with acceptable safety profile.

## Materials and Methods Materials

### Cell line, Drugs and Chemicals

The DAL tumor cell line was procured from National Centre for Cell Sciences, Pune, India and cultured in DMEM supplemented with 10% FCS and 1% antibiotics (penicillin and streptomycin). Standard anti-cancer drug Methotrexate was obtained from the IPCA laboratories (Mumbai, India). Emodin compound was synthesized and outsourced from Hi-Media, Drabkin’s solution from Nice Chemicals Pvt. Ltd. (Cochin, India). All the chemicals were procured from Merk.

### Animals

Swiss albino mice of 20 to 25g weight were purchased from Kerala Veterinary and Animal Sciences University and house in polyacrylic cages at 6 mice per cage with rice hulls as bedding material. The mice were maintained in ventilated cages, fed with normal mouse chow (Sai Feeds, Mumbai, India) and water ad libitum. The mice were maintained under standard laboratory conditions at room temperature 25°C ± 2°C with proper ventilation and a proper light-dark cycle was maintained for every 12 hrs. All the animal experiments are performed based on the rules and regulations assigned by Committee for the Purpose of Control and Supervision of Experiments on Animals (CPCSEA) guidelines, Government of India. Ethical Approval are obtained from Institutional Animal Ethics Committee, of Vignan’s Foundation for Science, Technology, and Research, and Karunya University before commencing the work.

### Cell culture

The DAL tumor cells were thawed once they were received from the supplier and washed with sterile chilled 1× PBS to get rid of DMSO and other stabilizers. The cells quality was checked before seeding them for culturing. Once the cells quality was determined, they were seeded in T-25 flasks in a total of 5 mL of complete DMEM with appropriate antibiotics. The cells were cultured and passed several times and frozen for further use.

### Anti-lymphoma study

Lymphoma was induced in 30 mice by administering the DAL cells through Intraperitoneal Injection route (1.5×10^6^ cells/mice) and monitored for 15 days **(Figure 1)**. The lymphoma induced mice and 6 normal mice without any tumor administration were further categorized into five groups each receiving different treatments as follows:

**Figure 1.**
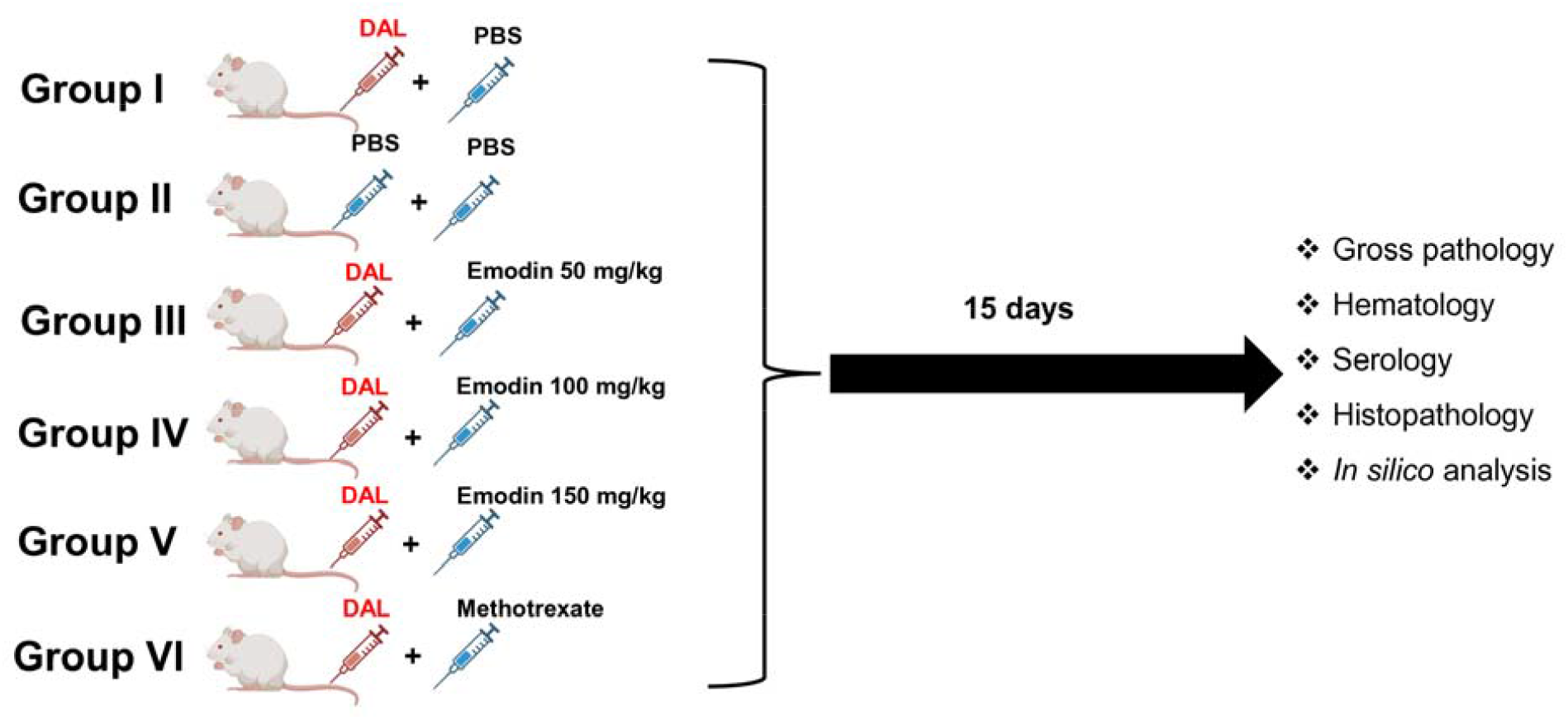
Schematic representation of the current study. Swiss Albino mice were divided into six groups (n=6). Group I-DAL+PBS (control); Group II-PBS+PBS (normal); Group III-DAL+50 mg/kg of emodin; Group IV-DAL+100 mg/kg of emodin; Group V-DAL+150 mg/kg of emodin; Group VI-DAL+methotrexate.

a. Group I-DAL+PBS.
b. Group II-normal.
c. Group III-DAL+50 mg/kg of Emodin.
d. Group IV-DAL+100 mg/kg of Emodin.
e. Group V-DAL+150 mg/kg of Emodin.
f. Group VI-DAL+Methotrexate.

The treatment was followed for 15 days following the previous studies (S Jagadish Kumar, 2016). After the treatment schedule, all the mice were sacrificed humanely and the *in vivo* therapeutic efficacy of Emodin was studied and compared with the controls, normal and Methotrexate treated mice. The *in vivo* therapeutic efficacy was studied by measuring the tumor, total body measurements, blood parameters, serum profile and histopathology.

### Tumor and total body measurements

After 15 days of the treatment schedule, all the mice were sacrificed, total body weight and tumor weight and volumes were measured in the control and treatment groups as part of studying the anti-tumor potential of emodin. The total body weight of each mouse in all the five groups was measured and noted after the treatment schedule. To measure the tumor weight and volume, the ascitic fluid was collected and measured its weight and volume in each mouse.

### Hematological parameters

Similarly, the blood was collected from each mouse in a heparin coated tube to prevent the coagulation of blood. The total hemoglobin content, RBC and WBC population were determined in each sample following the standard procedures according to our previous study (S Jagadish Kumar, 2016).

### Serology

An aliquot of blood was collected in a separate tube while collecting the blood. The blood was allowed to clot and centrifuged at 2000 rpm for 20 mins at 4^°^C to separate the serum from the other blood components. The serum was used to measure the Alkaline Phosphatase (ALP) and Aspartate amino transferase (AST) levels in each mouse from all the groups. The serum levels of ALP and AST were determined using the kits procured from Sigma-Aldrich.

### Histopathology

The tissue pathology of liver sections was studied in a representative mouse from control, normal and treated groups. The sections were sent to a local pathologist to cut and stain the sections with Hematoxylin and eosin and the slides were observed under a light microscope at 40X magnification.

### Drug likeliness

The drug likeliness properties of emodin were assessed *in-silico* based on the Lipinski’s rule of 5 and these properties of emodin were compared the standard drug Methotrexate. These properties were determined using the Swiss ADME tool (http://www.swissadme.ch/) based on the structure of molecule obtained from PubChem database in SMILES format.

### Statistical analysis

The experimental readings were represented in mean+SD and all the statistical analysis was performed through one-way ANOVA using GraphPad prism version 10.3.1. The *P* values (i.e., \**P* < 0.05, \*\**P* < 0.01, \*\*\**P* < 0.001, \*\*\*\**P*<0.0001) were considered as significant.

## Results

### Gross pathology

Increase in the total body weight is considered as a significant hallmark of lymphoma as the lymphocytes continuously proliferate. As a result, the abdomen was distended in the tumor induced mice. The treatment of tumor induced mice with a standard drug or emodin has shown a significant effect on the total body weight in the tumor induced mice. Morphologically, the treatment has blunted the enlargement of abdomen in the mice thereby decreasing the overall body weight **(Figure 2A)**. Similarly, there is a significant decrease in the overall weight **(Figure 2B)**, tumor volume and weights **(Figure 2C)** in methotrexate emodin treatment groups at all the three concentrations.

**Figure 2.**
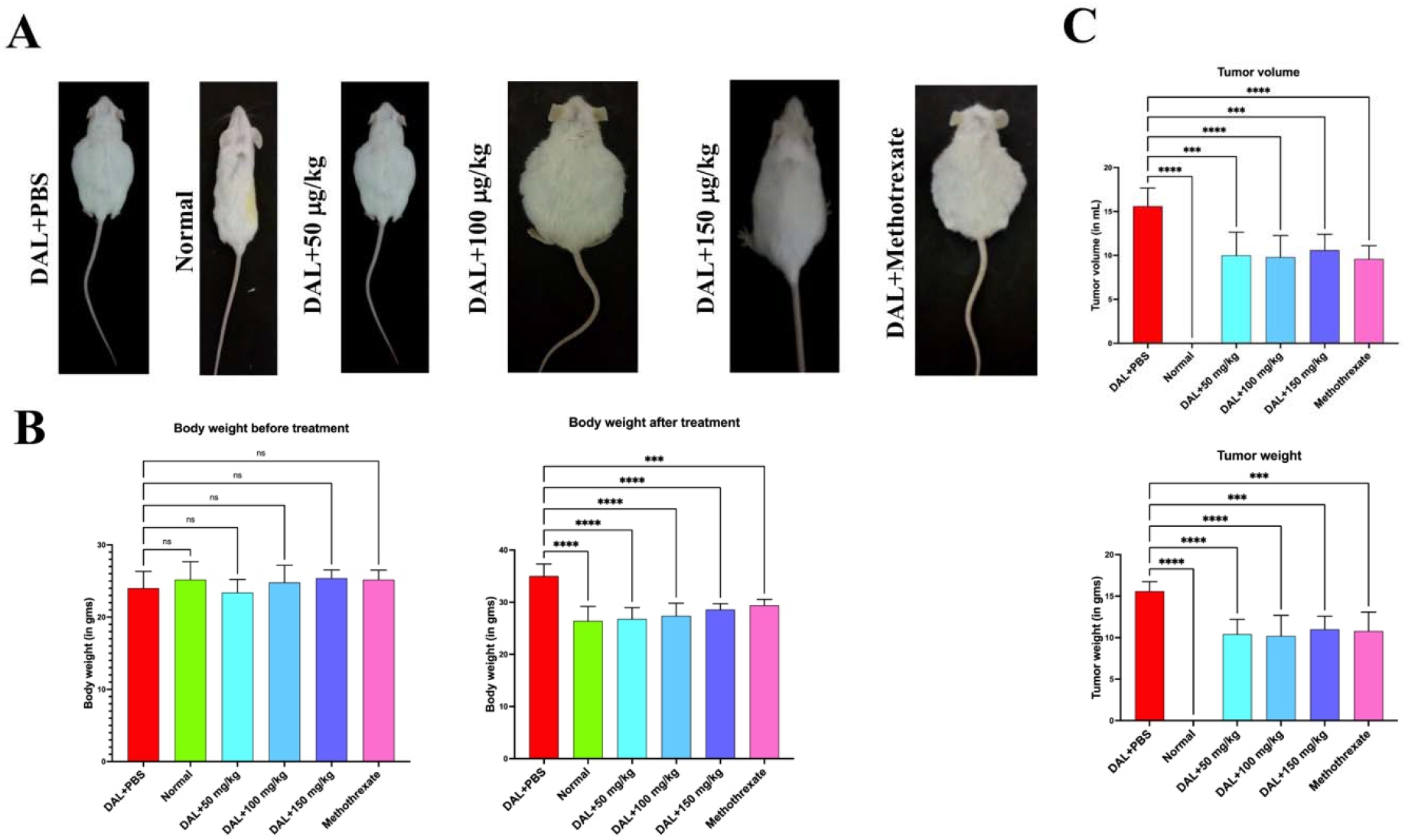
Therapeutic effect of emodin and methotrexate on the gross pathology of DAL induced mice. **A**. Emodin and methotrexate treatment has reduced the swelling in DAL induced mice; **B**. Reduced the body weights (50 mg/kg of emodin [P<0.0001]; 100 mg/kg of emodin [P<0.0001]; 150 mg/kg of emodin [P<0.0001]; and methotrexate [P=0.0003]); **C**. Tumor volume (50 mg/kg of emodin [P=0.0001]; 100 mg/kg of emodin [P<0.0001]; 150 mg/kg of emodin [P=0.0005]; and methotrexate [P<0.0001]) and weights (50 mg/kg of emodin [P<0.0001]; 100 mg/kg of emodin [P<0.0001]; 150 mg/kg of emodin [P=0.0004]; and methotrexate [P=0.0002]) in treated groups.

### Hematology

The total RBC population, hemoglobin content and WBC population were estimated after sacrifice of all the animals. The RBC count is reduced in the tumor group as compared to the normal group. While the treatment with methotrexate or emodin at 100 mg/kg has shown a positive effect on the RBC population by significantly increasing the cell count but not in the other groups **(Figure 3A)**. The similar trend is seen in the case of hemoglobin content among all the groups **(Figure 3B)**. The percentage of hemoglobin content is significantly increased in all the treatment groups as compared to the control ones. On the other hand, the emodin treatment has shown its therapeutic effect by reducing the total WBC population at all the concentrations like the methotrexate treatment **(Figure 3C)**, which highlights the relevance of emodin as a potent anti-lymphoma drug by reducing the total lymphocytes/WBC population.

**Figure 3.**
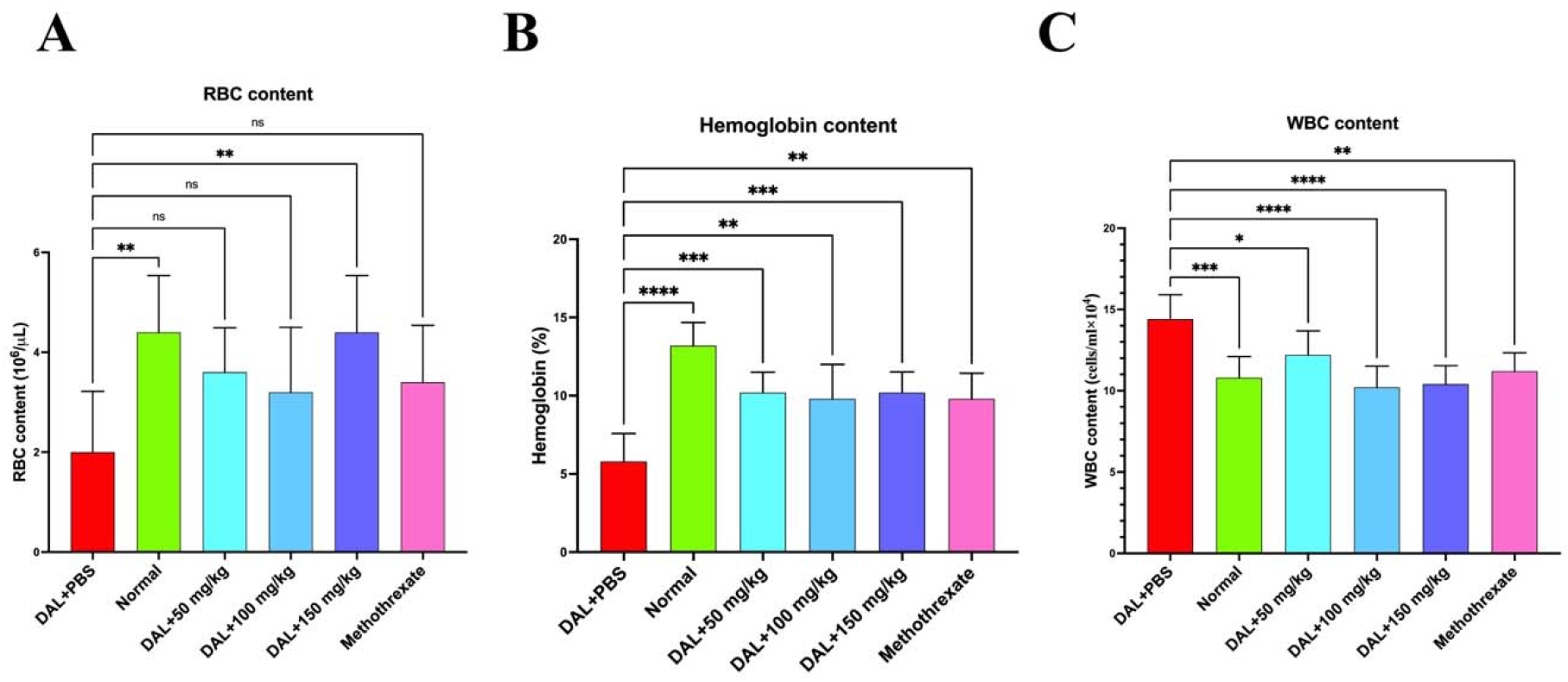
Therapeutic effect of emodin and methotrexate on the hematological parameters of DAL induced mice. **A**. RBC count in 50 mg/kg of emodin [P=0.0838]; 100 mg/kg of emodin [P=0.2668]; 150 mg/kg of emodin [P=0.0046]; and methotrexate [P=0.1542]), **B**. Hemoglobin content in 50 mg/kg of emodin [P= 0.0003]; 100 mg/kg of emodin [P= 0.0010]; 150 mg/kg of emodin [P= 0.0003]; and methotrexate [P= 0.0010]), **C**. WBC count in 50 mg/kg of emodin [P= 0.0295]; 100 mg/kg of emodin [P <0.0001]; 150 mg/kg of emodin [P <0.0001]; and methotrexate [P= 0.0010]).

### Serology

The serum biomarkers of liver such as alanine phosphatase (ALP) and aspartate aminotransferase (AST) are used to monitor the effectiveness of anti-lymphoma chemotherapeutic drugs. We also studied the effect of emodin treatment on these markers along with the standard methotrexate treatment. The ALP and AST levels are very high in the tumor groups than the normal mice **(Figure 4A&B)**. The levels of these two markers were also greatly reduced in all the treatment groups and reached the normal level **(Figure 4A&B)**.

**Figure 4.**
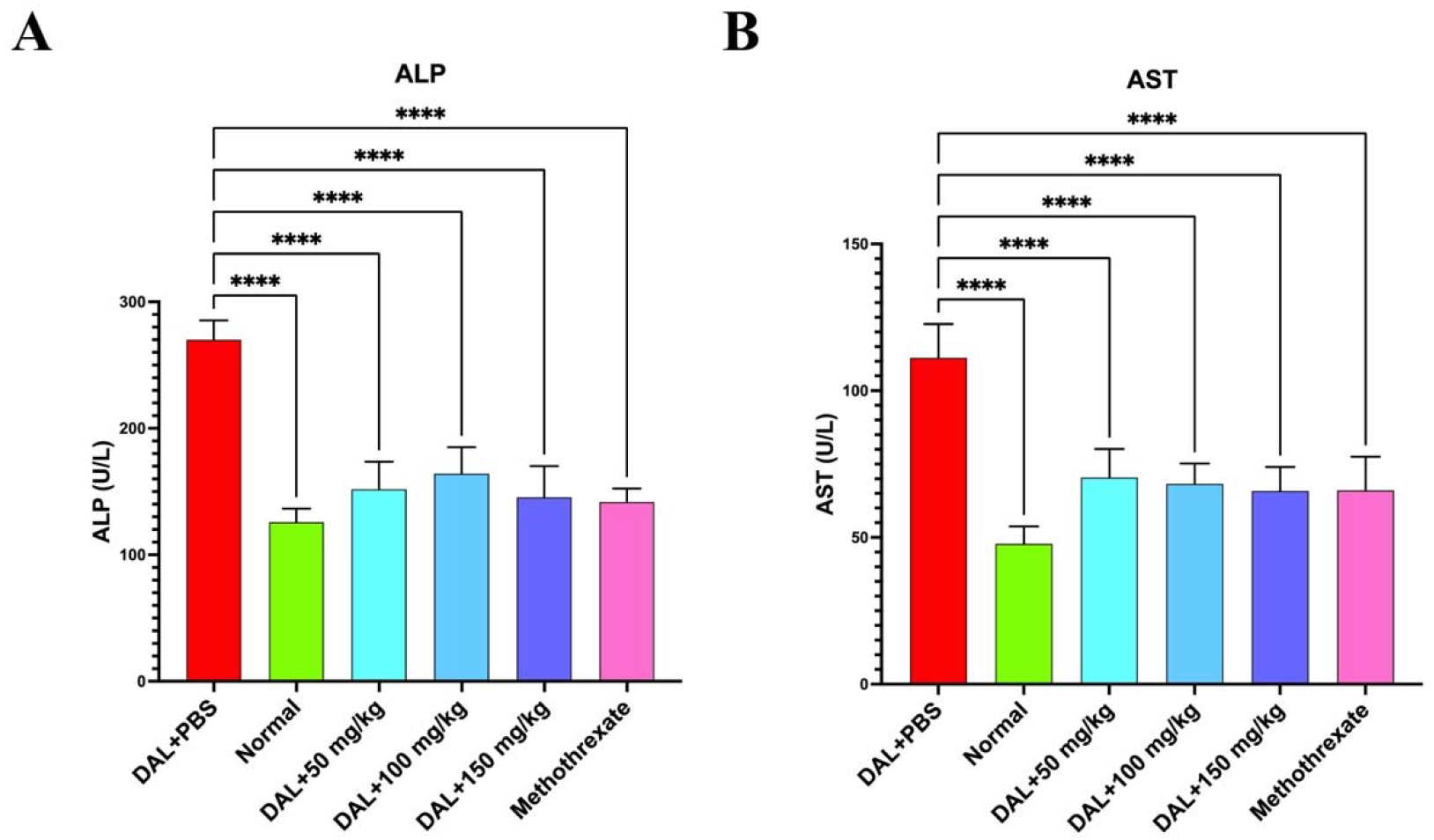
Therapeutic effect of emodin and methotrexate on the serum biomarkers in DAL induced mice. **A**. ALP levels in 50 mg/kg of emodin [P<0.0001]; 100 mg/kg of emodin [P<0.0001]; 150 mg/kg of emodin [P<0.0001]; and methotrexate [P<0.0001]), **B**. AST levels in 50 mg/kg of emodin [P<0.0001]; 100 mg/kg of emodin [P<0.0001]; 150 mg/kg of emodin [P<0.0001]; and methotrexate [P<0.0001]).

### Histopathology

The liver pathology was studied at the end of experiments by staining the liver sections with H&E stain. Histopathological observations have shown that the liver section of normal mice that received PBS alone has normal lobular architecture with preserved hepatocyte morphology. Whereas the tissue sections from DAL tumor bearing mice show that the portal areas are surrounded with severe fibrosis and massive inflammation **(Figure 5A-B)**. While this pathological condition is found to be reduced in all the treatment groups with the normal hepatocellular architecture evidencing the anti-tumor activity of emodin **(Figure 5C-F)**.

**Figure 5.**
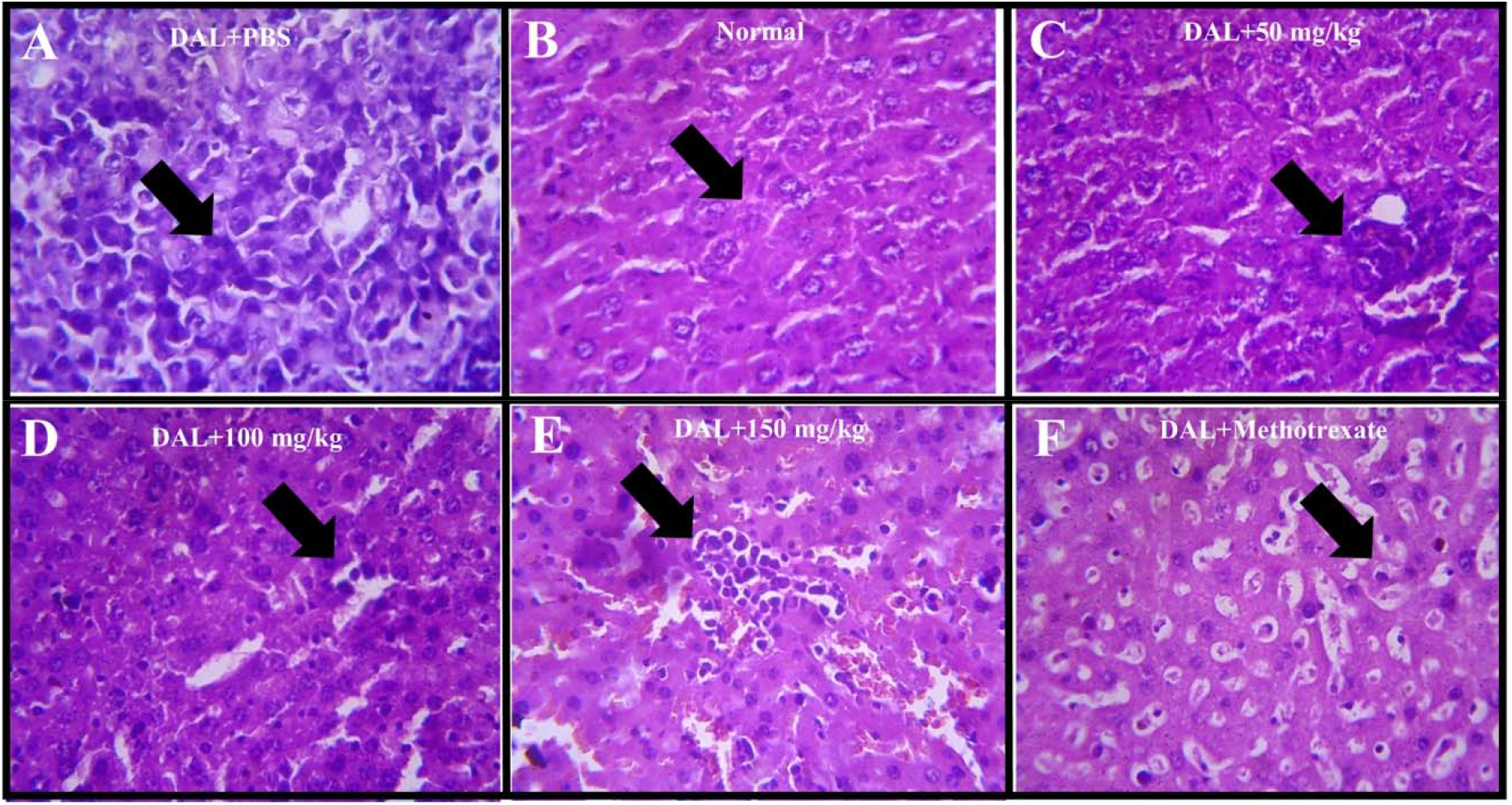
Histopathological examination of liver sections from the representative mouse in. **A**. DAL+PBS (control), **B**. PBS+PBS (normal), **C**. 50 mg/kg of emodin, **D**. 100 mg/kg of emodin, **E**. 150 mg/kg of emodin, **F**. and methotrexate treated groups.

### Drug likeliness

The drug likeliness properties of emodin compound were assessed based on the Lipinski’s rule of five-molecular weight, hydrogen bond donors, acceptors and LogP values. Based on the *in-silico* analysis, it is evident that the emodin compound has satisfied all these criteria with a molecular weight of below 500 g/mol, less than 5 H-donors and 10 H-bond acceptors and with a LogP value of less than 5 **(Figure 6)**. All these values were compared with the standard drug methotrexate and the emodin was found to be a better drug candidate to treat lymphoma.

**Figure 6.**
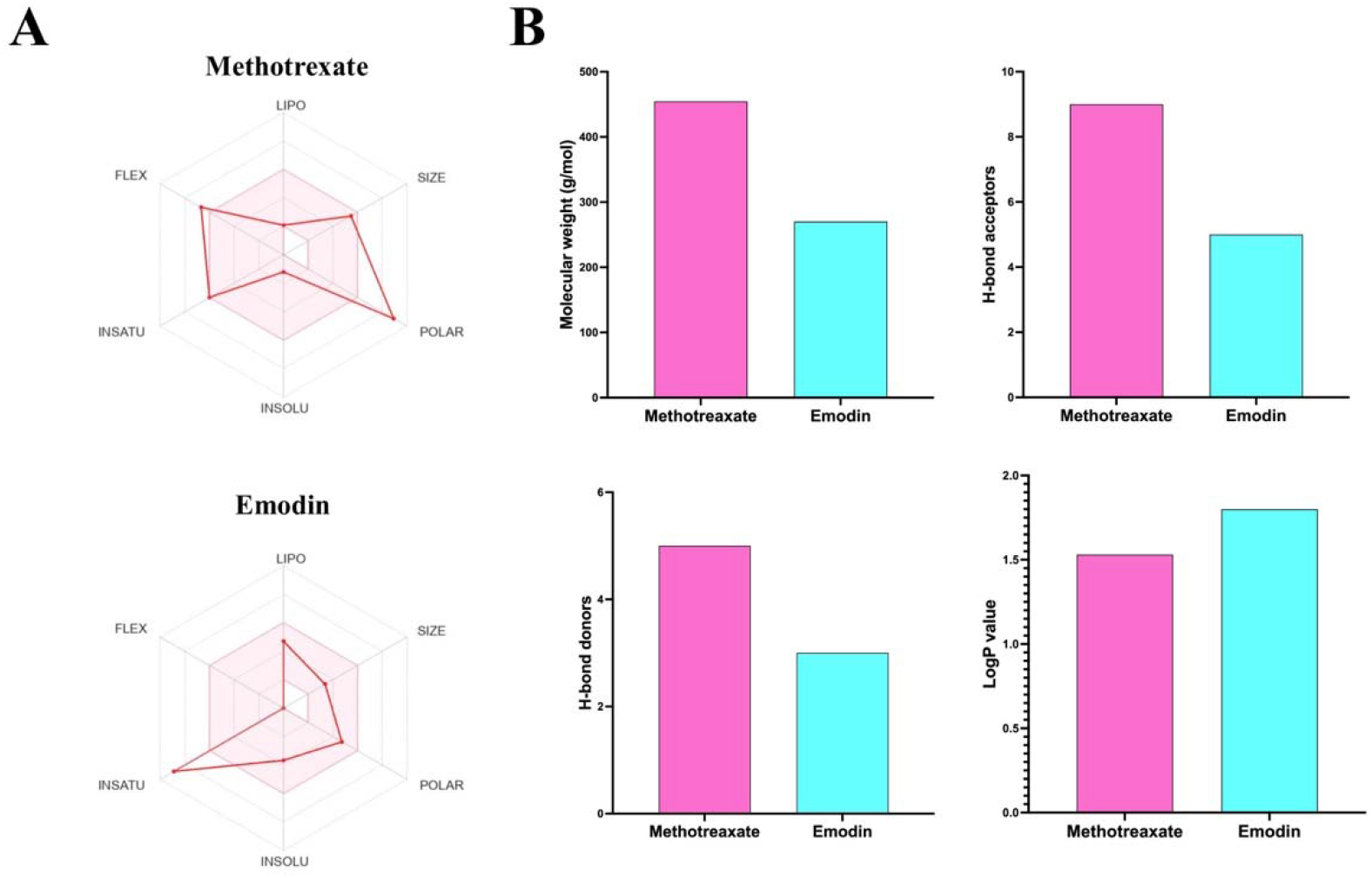
Drug likeliness profile of emodin and methotrexate compounds. **A**. Swiss MADE map of emodin, **B**. Swiss ADME map of methotrexate, **C**. Lipinski’s rule of 5 parameters of emodin and methotrexate.

## Discussion

In the present study we elucidated the anti-lymphoma activity of a natural compound emodin in the Swiss albino mice model induced with the DAL ascites. Our findings highlight the potential of emodin in treating lymphoma in DAL induced murine model where the treatment significantly reduced the tumor burden which is studied through several readouts. Emodin is a natural anthraquinone molecule derived from various plant sources (Stompor-Goracy, 2021). It has been studied for its anti-cancer activity primarily because of its ability to induce apoptosis in the cancer cells and eliminate them (Zhang et al., 2020).

Our results indicate that the tumor inhibiting nature of emodin through the reduced body weight, tumor weight and volumes in the treated mice. Being an immunosuppressive agent (Qiu et al., 2020), the main aspect of emodin action against the lymphoma lies in its ability to reduce the WBC/lymphocyte population. We observed this nature of reducing or controlling the WBC population by emodin in our later results in the serology experiments.

Another important characteristic of emodin’s action is through its anti-inflammatory properties which is evident from our serology and liver histopathological observations (Cheng et al., 2022; Han et al., 2015; Li et al., 2020; Saha & Ahmad, 2024). The liver tissue is characterized by the presence of immune cell infiltration and increased tissue necrosis with an increase in the liver functional enzymes such as ALP and AST which promotes the growth of tumor and metastasis. We found that the emodin treatment has reduced the immune cell infiltration with remarkable reduction in the liver damage biomarkers ALP and AST in the emodin treated mice. This highlight the anti-inflammatory nature of emodin which might be attributed to the inhibition of nuclear factor kappa B pathway which is chronically activated in malignancies and contribute to severe inflammation and tissue damage (Zhu et al., 2023). Emodin not only reduced the inflammation but also create a less conductive environment for tumor progression. Furthermore, the *in-silico* analysis also has shown that the drug likeliness of emodin as a potential anti-lymphoma drug.

While our study highlighted the anti-lymphoma activity of emodin, there are several limitations as well. Firstly, the DAL model mimics the human lymphoma pathology but does not completely replicate it. Further studies on the efficacy of emodin in more murine and other higher animal models would appreciate the anti-lymphoma nature of this compound. Additionally, since the emodin can modulate multiple pathways in the cancer pathogenesis, more studies on its ability to modulate those pathways in controlling the lymphoma would be highly appreciated. Furthermore, in depth investigation of the pharmacokinetics, pharmacodynamics and metabolism of emodin in the body will be critical in standardizing the optimal dosage for clinical application.

## Conclusion

In this study we investigated the anti-lymphoma activity of emodin in the DAL-induced Swiss Albino mice. The mice bearing lymphoma has shown an improved therapeutic outcome with emodin treatment at different doses of 50 mg/kg, 100 mg/kg and 150 mg/kg in terms of the pathology and serological biomarkers of lymphoma respectively. These findings were compared along with a standard drug methotrexate treatment. The emodin has shown a better or considerable therapeutic activity in controlling the severity of lymphoma in comparison with the methotrexate. Our results show that the potential of emodin in treating lymphoma. However, further studies need to be carried out before translating the molecule as a drug candidate.

## Acknowledgements

The authors are thankful to the management of Vignan’s Foundation for Science, Technology, and Research and Karunya Universities.

## Author contributions

JKS planned and performed the experiments. HBK analyzed data, prepared the figures, wrote and revised the manuscript. JJ designed the study and supplied the necessary reagents.

## Availability of data and materials

The authors declare that all the data supporting the findings of this study are available within the paper.

## Declarations

## Ethical approval and consent to participate

All the experiments carried out in the current study were according to the guidelines from Institutional Animal Ethics Committee, of Vignan’s Foundation for Science, Technology, and Research (Reg. No.: 2046/ PO/ReBi/S/18/CPCSEA), Guntur, Andhra Pradesh, India.

